# The maternal brain is more flexible and responsive at rest: effective connectivity of the parental caregiving network in postpartum mothers

**DOI:** 10.1101/2022.09.26.509524

**Authors:** Edwina R Orchard, Katharina Voigt, Sidhant Chopra, Tribikram Thapa, Phillip GD Ward, Gary F Egan, Sharna D Jamadar

## Abstract

The field of neuroscience has largely overlooked the impact of motherhood on brain function outside the context of responses to infant stimuli. Here, we apply spectral dynamic causal modelling (spDCM) to resting-state fMRI data to investigate differences in brain function between a group of 40 first-time mothers at one-year postpartum and 39 age- and education-matched women who have never been pregnant. Using spDCM, we investigate the directionality (top-down vs bottom-up) and valence (inhibition vs excitation) of functional connections between six key brain regions implicated in motherhood: the dorsomedial prefrontal cortex, ventromedial prefrontal cortex, posterior cingulate cortex, parahippocampal gyrus, amygdala, and nucleus accumbens. We show a selective modulation of inhibitory pathways related to differences between (1) mothers and non-mothers, (2) the interactions between group and cognitive performance and (3) group and social cognition, and (4) differences related to maternal caregiving behaviour. Across analyses, we show consistent disinhibition between cognitive and affective regions suggesting more efficient, flexible, and responsive behaviour, subserving cognitive performance, social cognition, and maternal caregiving. Together our results support the interpretation of these key regions as constituting a parental caregiving network. The nucleus accumbens and the parahippocampal gyrus emerging as ‘hub’ regions of this network, highlighting the global importance of the affective limbic network for maternal caregiving, social cognition, and cognitive performance in the postpartum period.

---

The transition to motherhood is associated with both structural and functional neural plasticity across pregnancy and the postpartum period^1-7^, with brain changes that are long-lasting^8,9^ and may persist across the lifespan^10-14^. Studies of brain function in motherhood have largely focused on maternal neural responses to infant stimuli^15-17^, with several brain regions showing consistent activation in response to infant cues, including visual, auditory, tactile, and olfactory signals. As these regions have been commonly found in functional neuroimaging studies of motherhood, they have been proposed to form a putative ‘parental caregiving brain network’^17^. Regional activation of the parental caregiving network in response to infant stimuli is associated with better maternal attachment and positive caregiving behaviours, whereas aberrant activation of these regions is associated with postpartum depression^18-21^, substance use^22,23^, and trauma^24,25^. This suggests that activation of the parental caregiving network may support the initiation and maintenance of complex and sensitive maternal behaviours^17^. While each region of the putative parental caregiving network is consistently implicated in maternal behaviour, the network itself has not been theoretically formalised, and it is unknown how regions within the network influence each other to facilitate maternal caregiving.

The field of parental neuroscience has largely overlooked the impact of motherhood on brain function outside the context of responses to infant stimuli (e.g., during a resting-state). Resting-state connectivity is particularly interesting, as it is considered an index of ‘intrinsic’ brain function, and provides the neural framework for the individual’s full range of extrinsic responses^26,27^. Changes in resting-state brain function associated with motherhood may uncover global, generalised changes in brain function, with broader impacts for maternal behaviour, beyond responses to infant stimuli. A small number of seed-based resting-state connectivity studies have measured connectivity in *a priori* defined networks, and these studies have shown differences between mothers and non-mothers in six key regions of the putative parental caregiving network: the dorsomedial prefrontal cortex, ventromedial prefrontal cortex, parahippocampal gyrus, posterior cingulate cortex, amygdala, and nucleus accumbens^3,18,22,24,28-30^ (Figure 1a). Resting-state functional connectivity between the amygdala, nucleus accumbens, and dorsomedial prefrontal cortex are related to positive maternal care. Increased connectivity between these regions is associated with improved ability to scaffold appropriate mother-child interactions^3^ and higher maternal-child synchrony (the coordination of gaze, touch and vocalisations between mother and child)^24^. Additionally, higher-order association areas, including the dorsomedial prefrontal cortex and posterior cingulate cortex, show a positive relationship between functional connectivity and cognitive performance in mothers^28,29^, as illustrated in Figure 1b.

**Figure 1:**
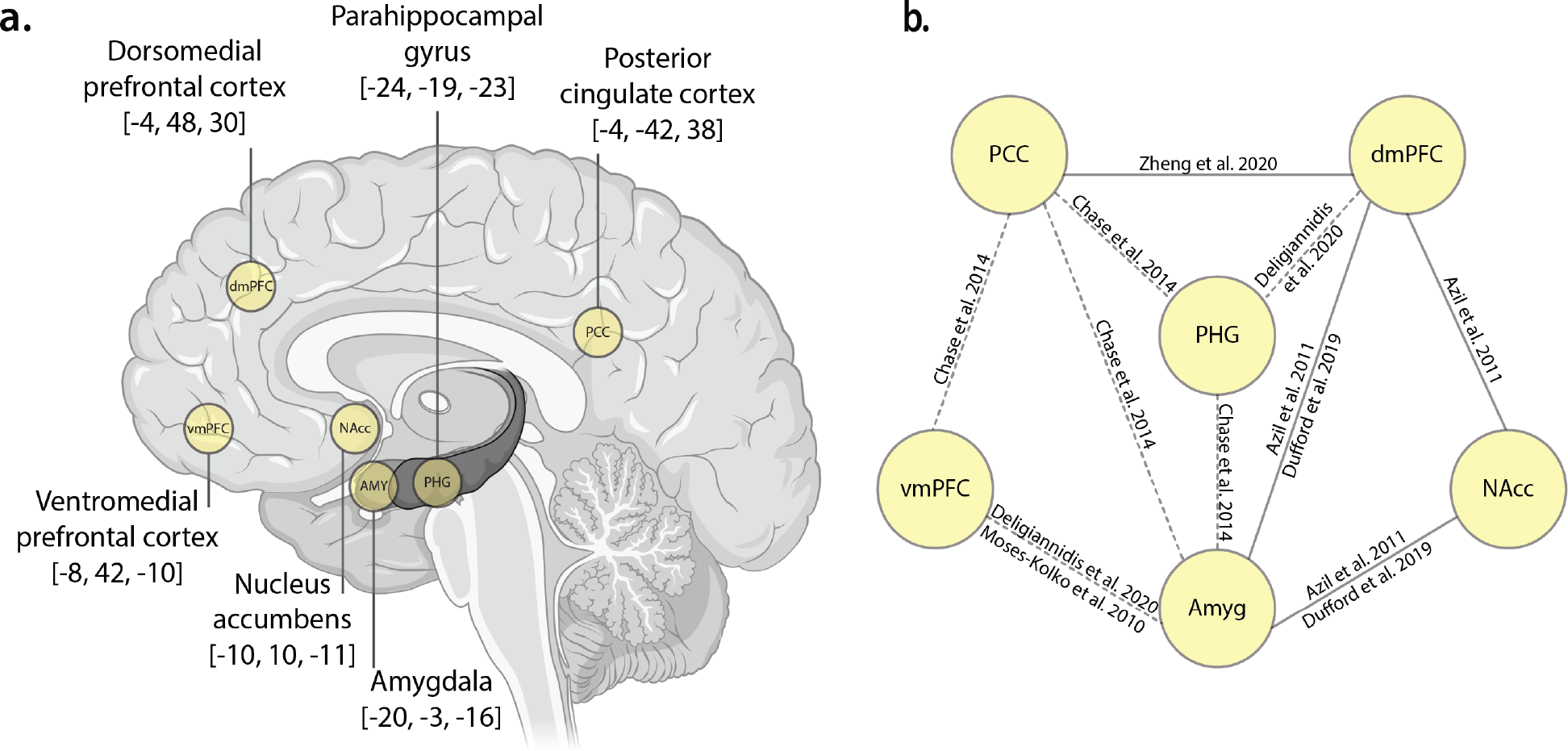
a.) Regions of interest. Six left-hemisphere Regions of Interest (ROIs), with MNI coordinates (x,y,z). Brain image created with BioRender. b.) Previously reported differences in functional connectivity. Schematic diagram showing the extant resting state functional connectivity literature in motherhood for selected regions of the parental caregiving brain network. Dashed lines represent connections which differ between mothers with and without postpartum depression. These known connections were used to formulate the study hypotheses. Abbreviations: PCC, posterior cingulate cortex; vmPFC, ventromedial prefrontal cortex; dmPFC, dorsomedial prefrontal cortex; PHG, parahippocampal gyrus; Amyg, amygdala; NAcc, nucleus accumbens.

Beyond this, much of what is known about maternal brain function at rest is from studies comparing mothers with and without postpartum depression. Mothers with postpartum depression show aberrant connectivity between the posterior cingulate cortex, parahippocampal gyrus, amygdala, ventromedial prefrontal cortex, and dorsomedial prefrontal cortex^18,22,30^, suggesting these regions are important for maternal caregiving, and are sensitive to, or involved in, postpartum mood disorders. Very little is known about the differences in resting state brain function between non-mothers and mothers without mood disorders.

The investigation of maternal brain function outside of the context of postpartum depression, and reactivity to infant stimuli, is important in order to characterise the ‘normative’ functional neural adaptations to motherhood. Importantly, while the parental caregiving network has been proposed to account for commonly activated regions found in neuroimaging studies of motherhood, no study has explicitly tested the network outside of infant-centric stimuli. Furthermore, an important step towards understanding the functional adaptations to motherhood is to examine the *direction* and *valence* of information flow throughout the parental caregiving brain network. Functional connectivity analyses indicate coherence of timeseries between spatially distinct regions, but do not provide information on the directionality of the connections. By contrast, *effective connectivity* analyses indicate the direction of connections (e.g., whether they represent top-down or bottom-up information flow) as well as the valence of the connections (whether they are excitatory or inhibitory). Understanding the dynamics of information flow within the parental caregiving brain network is an important step to understanding the functional adaptations to motherhood in the early postpartum period.

We investigated differences between mothers and non-mothers in effective connectivity between regions of the putative parental caregiving network, using spectral dynamic causal modelling (spDCM)^31^. SpDCM can map specific circuits that influence functional brain differences between mothers and non-mothers and enables the investigation of effective brain connectivity in maternal caregiving behaviour, cognitive performance, and social cognition.

Six regions of the parental caregiving network^17^ consistently emerge across functional connectivity studies in motherhood, showing relationships with maternal caregiving, cognition, and postpartum depression. Other regions are also implicated, but less consistently^3,18,22,24,28-30^. Informed by this understanding, we tested four hypotheses. First, to map the connectivity of the parental caregiving network, we hypothesised that mothers and non-mothers would show differences in effective connectivity between the dorsal and ventral medial prefrontal cortices, posterior cingulate cortex, parahippocampal gyrus, amygdala, and nucleus accumbens. Second and third, we hypothesised that an increased influence of higher-order regions, including dorsomedial prefrontal cortex, ventromedial prefrontal cortex, posterior cingulate cortex, and parahippocampal gyrus^32-34^, would be related to both *cognitive performance* (verbal memory, working memory, and processing speed); and *social cognition* (empathy and theory of mind). Fourth, we hypothesise that *maternal caregiving* (attachment and self-efficacy) would be related to an increased influence of the amygdala and nucleus accumbens, consistent with functional connectivity studies of maternal caregiving behaviour^3,24^.

## Results

Mothers and non-mothers did not differ in age or education^35^. Compared to non-mothers, mothers had poorer sleep and higher depression. Mothers also scored higher than non-mothers on the wellbeing component, which combines depression, anxiety, and sleep, indicating poorer wellbeing in mothers. Mothers and non-mothers did not differ on the cognitive performance component. Mothers scored higher than non-mothers on the social cognition component, indicating better social cognition. The means and standard deviations for the cognition and social cognition variables within the principal components, and significance tests, are reported in Supplementary Information 3.

At the subject-level, diagnostic statistics ensured first-level model inversion had converged. The average variance explained across subject-level DCM inversion was very high (> 85, mothers: *M*=87.6, *SD*=1.6, range=82.4-91.4; non-mothers: *M*=87.9, *SD*=2.4, range =79.9-92.3). The results of each of the four models are shown in Figure 2. All significant connections in all four models were inhibitory connections.

**Figure 2:**
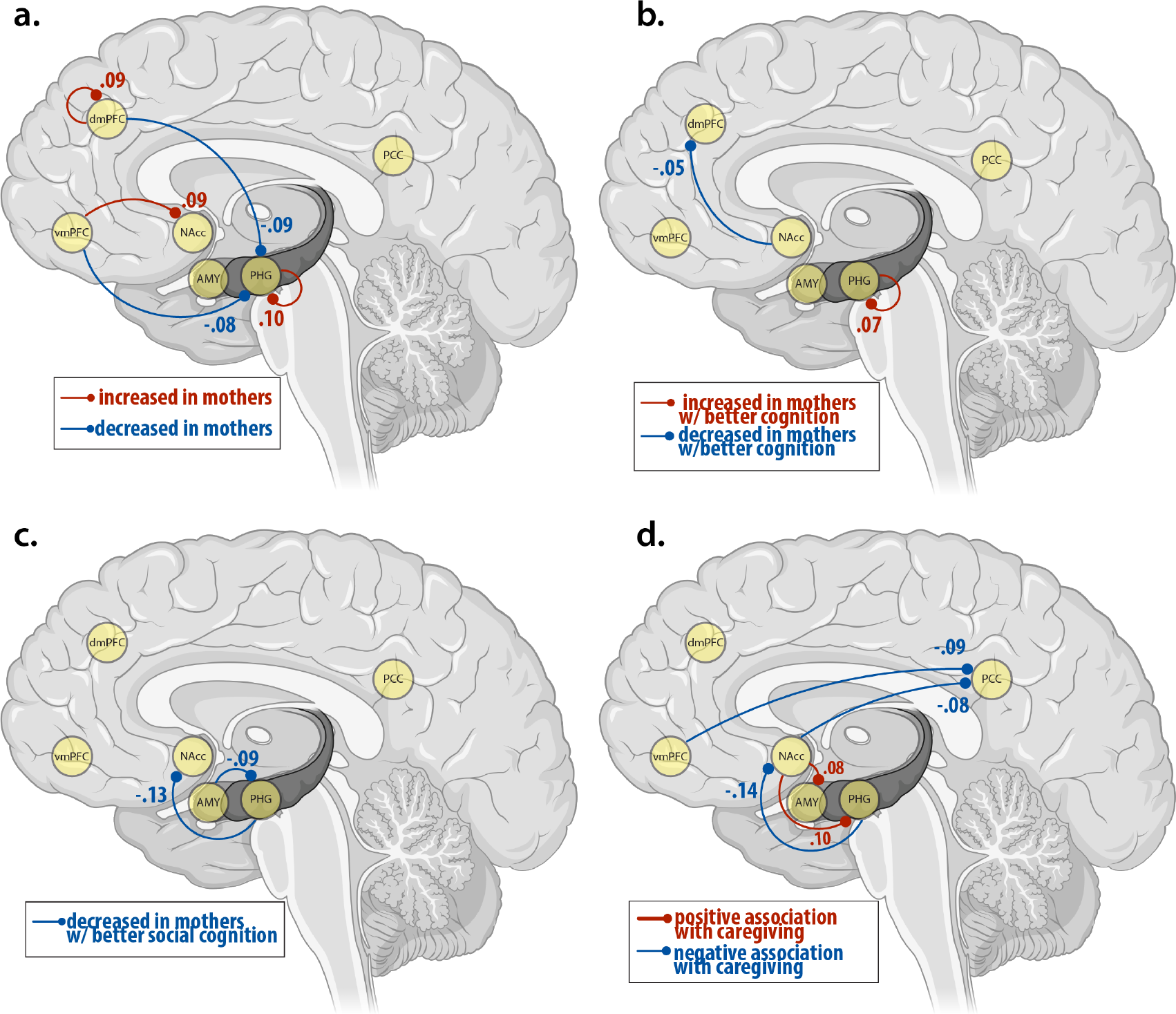
Schematic diagram showing spDCM results for (a.) the main effect of group (mothers vs non-mothers), (b.) the interaction effect of group-by-cognition, (c.) the interaction effect of group-by-social cognition, and (d.) the main effect of maternal caregiving in mothers only. All connections are inhibitory. Values represent effect sizes (see Table 2-5) in Hz. Blue lines depict decreased inhibition and red lines depict increased inhibition. Dashed lines represent connections that are attenuated when the model adjusts for wellbeing, and solid lines are connections which are present whether or not wellbeing is adjusted for. Abbreviations: PCC, posterior cingulate cortex; vmPFC, ventromedial prefrontal cortex; dmPFC, dorsomedial prefrontal cortex; PHG, parahippocampal gyrus; Amyg, amygdala; NAcc, nucleus accumbens.

**Table 1:** Demographic information for Mothers and Non-Mothers, including independent samples t-test comparisons between groups for age, education, sleep (Pittsburgh Sleep Quality Index), depression (Beck Depression Inventory), and anxiety (Beck Anxiety Index).

|  | Mothers (N=40) |  | Non-Mothers (N=39) |  | *t*-value | Cohen's *d* | *p*-value |
| --- | --- | --- | --- | --- | --- | --- | --- |
|  | Mean (SD) | Range | Mean (SD) | Range |  |  |  |
| Age (years) | 32.5 (3.3) | 25-40 | 31.3 (4.0) | 20-39 | 1.36 | 0.31 | .18 |
| Education (years) | 18.2 (2.5) | 13-24.5 | 17.6 (2.7) | 11-24 | 1.03 | 0.23 | .30 |
| Sleep (PSQI) | 7.3 (3.1) | 2-17 | 4.7 (2.6) | 0-10 | 4.17 | 0.94 | <.001 |
| Depression (BDI) | 8.6 (5.6) | 0-24 | 5.0 (4.3) | 0-17 | 3.20 | 0.72 | .002 |
| Anxiety (BAI) | 10.7 (8.6) | 0-32 | 8.6 (6.63) | 0-29 | 1.30 | 0.29 | .20 |

**Table 2:**
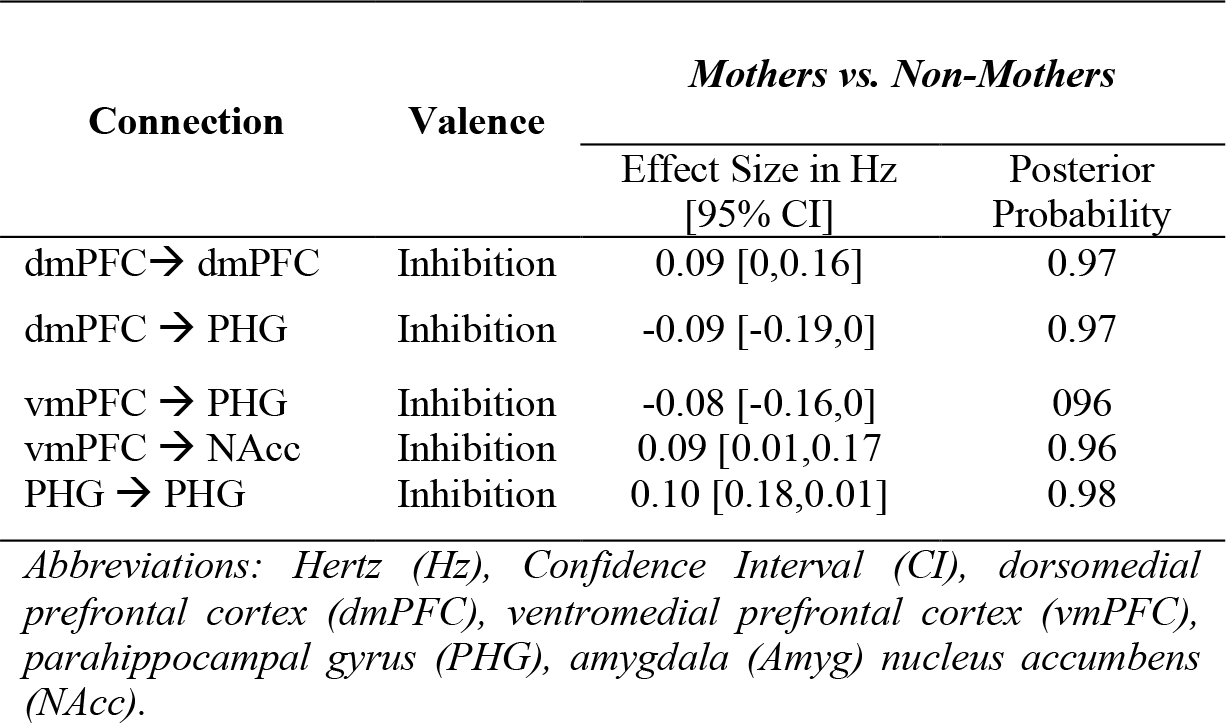
Effect size, valence and posterior probabilities for each connection showing a main effect of group (mothers>non-mothers)

| Connection | Valence | Mothers vs. Non-Mothers |  |
| --- | --- | --- | --- |
|  |  | Effect Size in Hz<br>[95% CI] | Posterior<br>Probability |
| dmPFC → dmPFC | Inhibition | 0.09 [0,0.16] | 0.97 |
| dmPFC → PHG | Inhibition | -0.09 [-0.19,0] | 0.97 |
| vmPFC → PHG | Inhibition | -0.08 [-0.16,0] | 0.96 |
| vmPFC → NAcc | Inhibition | 0.09 [0.01,0.17] | 0.96 |
| PHG → PHG | Inhibition | 0.10 [0.18,0.01] | 0.98 |

Abbreviations: Hertz (Hz), Confidence Interval (CI), dorsomedial prefrontal cortex (dmPFC), ventromedial prefrontal cortex (vmPFC), parahippocampal gyrus (PHG), amygdala (Amyg) nucleus accumbens (NAcc).

**Table 3:**
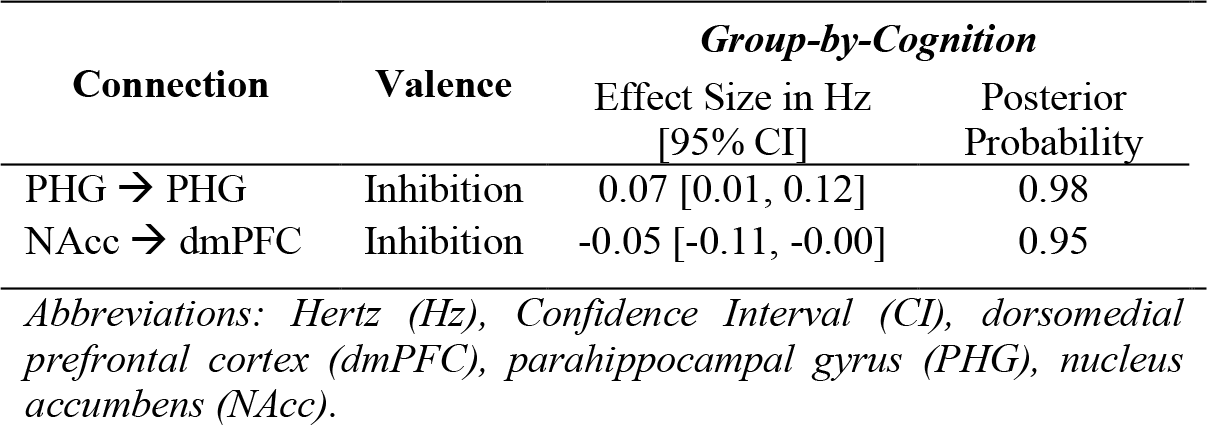
Effect size, valence and posterior probabilities for each connection showing an interaction effect of group-by-cognition

**Table 4:**
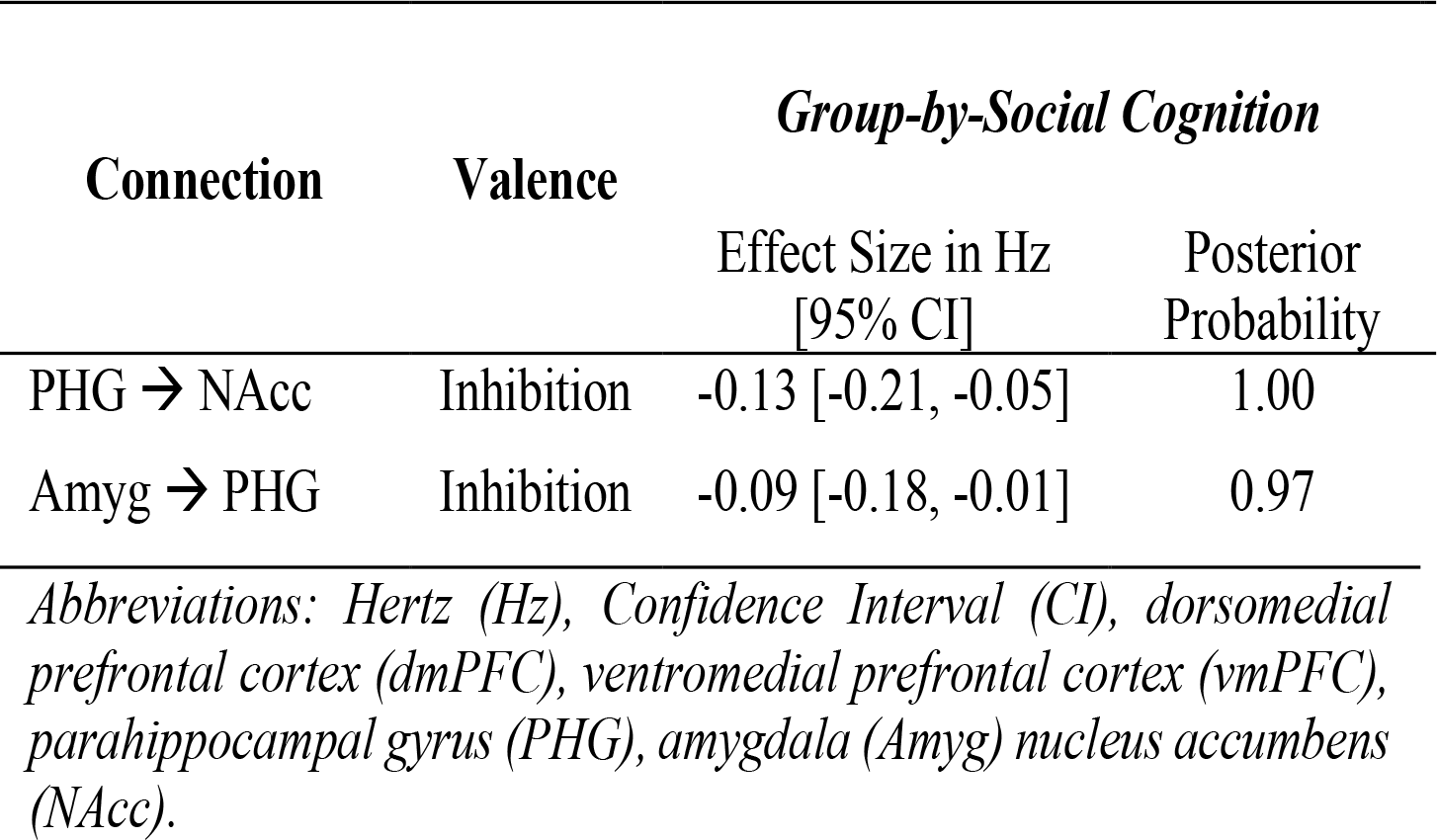
Effect size, valence and posterior probabilities for each connection showing an interaction effect of group-by-social cognition, with and without controlling for wellbeing

**Table 5:**
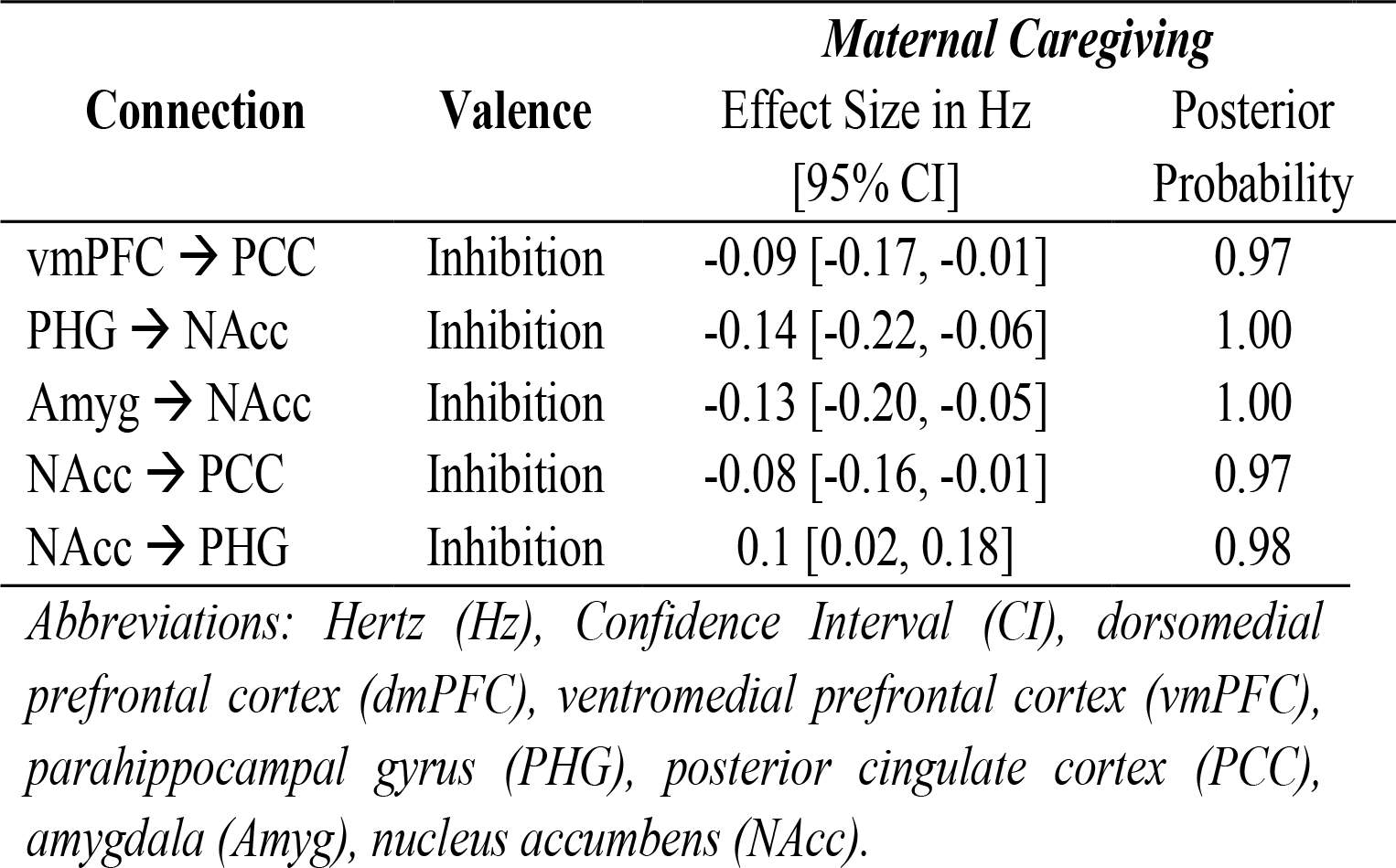
Main effect of maternal caregiving, with and without controlling for wellbeing

| Connection | Valence | Maternal Caregiving |  |
| --- | --- | --- | --- |
|  |  | Effect Size in Hz<br>[95% CI] | Posterior<br>Probability |
| vmPFC → PCC | Inhibition | -0.09 [-0.17, -0.01] | 0.97 |
| PHG → NAcc | Inhibition | -0.14 [-0.22, -0.06] | 1.00 |
| Amyg → NAcc | Inhibition | -0.13 [-0.20, -0.05] | 1.00 |
| NAcc → PCC | Inhibition | -0.08 [-0.16, -0.01] | 0.97 |
| NAcc → PHG | Inhibition | 0.1 [0.02, 0.18] | 0.98 |

Abbreviations: Hertz (Hz), Confidence Interval (CI), dorsomedial prefrontal cortex (dmPFC), ventromedial prefrontal cortex (vmPFC), parahippocampal gyrus (PHG), posterior cingulate cortex (PCC), amygdala (Amyg), nucleus accumbens (NAcc).

Compared to non-mothers, mothers showed decreased top-down inhibition of the parahippocampal gyrus from the dorsomedial prefrontal cortex, and the ventromedial prefrontal cortex, and increased top-down inhibition of the nucleus accumbens from the ventral medial prefrontal cortex (Table 2, Figure 2a). The dorsomedial prefrontal cortex and parahippocampal gyrus showed increased self-inhibition in mothers compared to non-mothers. Mothers with better cognitive performance showed decreased bottom-up inhibition from the nucleus accumbens to the dorsomedial prefrontal cortex, and increased self-inhibition of the parahippocampal gyrus (Table 3, Figure 2b). Mothers with better social cognition showed decreased top-down inhibition of the nucleus accumbens from the parahippocampal gyrus, and increased bottom-up inhibition from the amygdala to the parahippocampal gyrus (Table 4, Figure 2c). Within the mother group, higher maternal caregiving was associated with increased bottom-up inhibition from the nucleus accumbens to the parahippocampal gyrus, decreased inhibition from the nucleus accumbens and the ventromedial prefrontal cortex to the posterior cingulate cortex, and decreased inhibition to the nucleus accumbens from the amygdala and parahippocampal gyrus (Table 5, Figure 2d).

The nucleus accumbens and the parahippocampal gyrus emerged as ‘hub’ regions of the maternal network. Both the nucleus accumbens and the parahippocampal gyrus show significant connections in every model tested (main effect of group, main effect of maternal caregiving, interaction effect of group-by-cognition, and interaction effect of group-by-social cognition). When interpreting the results from our four models together, these two key regions show the highest nodal degree (number of connections), with 29% of all connections to or from the nucleus accumbens (9/31) and 26% of all connections to or from the parahippocampal gyrus (8/31). There were only three connections that did not include either the parahippocampal gyrus or the nucleus accumbens. For mothers versus non-mothers this included self inhibition of the dorsomedial prefrontal cortex; and in mothers with higher maternal caregiving, reduced inhibition of the posterior cingulate cortex from the ventromedial prefrontal cortex, and from the dorsomedial prefrontal cortex to the amygdala was observed. The strongest connections (effect size in Hz >0.1) between these hub regions were from the parahippocampal gyrus to the nucleus accumbens (-0.15 Hz), self-connections from the parahippocampal gyrus to itself (0.11 Hz), and from the nucleus accumbens to the parahippocampal gyrus (0.10 Hz). Other connections stronger than 0.10 Hz were from the amygdala to the nucleus accumbens (-0.13 Hz; maternal caregiving) and from the dorsomedial prefrontal cortex to the parahippocampal gyrus (-0.10 Hz; mothers vs non-mothers).

## Discussion

Understanding differences in brain connectivity at rest may uncover neural correlates of maternal care, and have broader implications for non-maternal behaviours, including cognition. We investigated the causal interactions between six key regions of the parental caregiving network (dorsomedial prefrontal cortex, ventromedial prefrontal cortex, posterior cingulate cortex, parahippocampal gyrus, amygdala, and nucleus accumbens), using spectral dynamic causal modelling (spDCM). Our results are the first experimental evidence to confirm this network is highly implicated in maternal caregiving. We showed that the dynamics of this network differ between non-mothers and mothers at one-year postpartum. Specifically, we show consistent patterns of disinhibition between cognitive and affective regions across all analyses, suggesting (1) a more responsive, flexible, and efficient emotion-regulation system in mothers compared to non-mothers, (2) increased selectivity for goal-directed information in mothers who perform better on cognitive tasks, (3) disinhibition between cognitive and affective mentalising networks related to both cognition and social cognition, and (4) a more selective affective limbic network in mothers with higher maternal caregiving. The results indicated that the nucleus accumbens and parahippocampal gyrus are ‘hub’ regions of the maternal caregiving brain network, as discussed in Supplementary Information 4.

### Effective connectivity of the parental caregiving network in mothers vs non-mothers

Relative to non-mothers, mothers showed increased self-inhibition of the dorsomedial prefrontal cortex and the parahippocampal gyrus. Self-connections regulate the excitatory and inhibitory balance of the network; controlling the *gain* (or excitatory/inhibitory balance) of the region to extrinsic connections^57^. In predictive coding theory, which is core to understanding the results of DCM analysis, the brain infers the cause of its sensory inputs by minimising prediction errors throughout the hierarchical cortical network^58,59^. Importantly, as self-connections regulate the gain or excitability of superficial pyramidal cells within a region, they also regulate the precision of the prediction error. Thus, the finding that mothers show increased self-inhibition of the dorsomedial prefrontal cortex and parahippocampal gyrus is consistent with the interpretation that motherhood is associated with increased network efficiency (e.g.,^60^) and more precise prediction error^59,61^ of the parental caregiving network relative to non-mothers. This improved precision for motherhood contrasts with pathologies like schizophrenia, which is associated with reduced self-connections and aberrant prediction error throughout the cortical hierarchy^59,61^.

Mothers also showed decreased top-down inhibition from the dorso- and ventro-medial prefrontal cortices to the parahippocampal gyrus; and from the ventromedial prefrontal cortex to nucleus accumbens, compared to non-mothers. Together, the dorsomedial prefrontal cortex, ventromedial prefrontal cortex, nucleus accumbens and parahippocampal gyrus form key regions of the *ventromedial emotion regulatory system*, which is implicated in early processing and encoding of the emotional significance of a stimulus^62^. This system is involved in early appraisal, encoding, and automatic regulation of emotion. The connection between dorsomedial prefrontal cortex to parahippocampal gyrus, and ventromedial prefrontal cortex to parahippocampal gyrus was reduced in mothers relative to non-mothers, indicating a less inhibited top-down circuit for mothers relative to non-mothers. A less inhibited network for mothers means implies the network is more easily excitable compared to non-mothers; in other words, less input is required to reach the ‘threshold’ for activation. Together with the increased self-connection of the dorsomedial prefrontal cortex and parahippocampal gyrus, this pattern of results indicates a more responsive, flexible and efficient emotion regulation system in mothers than non-mothers^62^.

### Differences in effective connectivity between mothers and non-mothers related to cognition

Behaviourally, mothers and non-mothers showed no significant differences in cognitive performance (verbal memory, working memory, and processing speed)^35^. The absence of group differences in cognitive performance may partly explain the low dimensionality of the effective connectivity network found in the group-by-cognition analysis.

Our spDCM results show that mothers with better cognitive performance showed decreased bottom-up inhibition from the nucleus accumbens to the dorsomedial prefrontal cortex, and increased self-inhibition of the parahippocampal gyrus. As noted above, self-inhibition within DCM reflects the excitation/inhibition balance of superficial pyramidal cells within a cortical region^57,59^. Increased self-inhibition of the parahippocampal gyrus in mothers with improved cognitive performance indicates reduced neural excitability and improved prediction error relative to non-mothers. The parahippocampal gyrus is involved in memory consolidation^63^ and mediates communication between the cortex and the hippocampus^64^. The parahippocampal gyrus may be considered a filter, or a gate, which selectively controls information transfer between the neocortex and the hippocampus through intrinsic inhibitory mechanisms^32,65^. Increased self-inhibition of the parahippocampal gyrus in mothers with better cognitive performance is consistent with memory consolidation and retrieval functions which require selective and precise integration of information to and from the hippocampus^32^. fMRI studies in healthy adults have shown the parahippocampal gyrus is a primary hub of the default mode network in the medial temporal pole, functionally coupled with cortical nodes of the default mode network, and mediates memory encoding at rest^66^. These findings suggest that mothers in the peripartum period may experience increased information selectivity by the parahippocampal gyrus, which may be related to cognitive performance in the peripartum period.

In addition, mothers with better cognitive performance showed decreased bottom-up inhibition of the dorsomedial prefrontal cortex from the nucleus accumbens. This connection was no longer significant once accounting for differences in wellbeing (Supplementary Information 2), indicating that this connection is affected by the individual’s level of depression, anxiety, and sleep quality. The dorsomedial prefrontal cortex and nucleus accumbens form core regions of the cortico-basal ganglia-thalamic network that underlies goal-directed motor behaviour to motivationally salient events^34,67,68^. The dorsomedial prefrontal cortex in particular is implicated in the control of actions within this network, and the input from the nucleus accumbens to the dorsomedial prefrontal cortex provides the emotional and motivational context to drive the appropriate motor response^34^. Importantly, the connection between the nucleus accumbens and dorsomedial prefrontal cortex is known to be mediated by sleep. The dorsomedial prefrontal cortex-nucleus accumbens network is known to be involved in consolidating remote, but not recent, memory^34^. A key part of memory consolidation, ‘replay’, occurs during sleep and neurons within these regions show spike patterns consistent with replay^69,70^. Furthermore, chronic blockage of the efferent medium spiny neurons of the nucleus accumbens alters the dorsomedial prefrontal cortex transcriptome, particularly gene sets associated with cognition, sleep, and movement^71^. Taken together, our results indicate that the bottom-up inhibitory connection from the nucleus accumbens to the dorsomedial prefrontal cortex may underlie wellbeing-mediated differences in cognition between mothers and non-mothers^35^.

### Differences in effective connectivity between mothers and non-mothers related to social cognition

Compared to non-mothers, mothers had significantly higher scores on measures of social cognition (theory of mind, and empathy; Supplementary Information 5). Mothers with higher social cognition scores had decreased inhibition from the amygdala to the parahippocampal gyrus, and from the parahippocampal gyrus to the nucleus accumbens, relative to non-mothers. The nucleus accumbens-amygdala-parahippocampal gyrus network forms the *affective limbic network* that is implicated in social connectedness, and its converse, loneliness^72^. This network overlaps with the subcortical components of the ventromedial emotion regulatory system, which is implicated in early appraisal, encoding, and automatic regulation of emotion^62^. The limbic network is particularly involved in affective aspects of theory of mind, pertaining to making inferences about emotions^33^, as well as assigning motivational value to goals on the basis of internal states and environmental stimuli^73^. Within this network, the amygdala and hippocampus establish the motivational value of goals, the prefrontal cortex encodes potential action outcomes, and the nucleus accumbens integrates different sources of values to guide the selection of appropriate behaviour^73^. That mothers with better social cognition showed decreased inhibition within this network indicates that the network is relatively disinhibited relative to non-mothers. In other words, the network can be more responsive, i.e., requires lower levels of input to activate, in mothers with improved social cognition. Consistent with our other analyses, mothers show a more responsive limbic network which supports their responsiveness to affective stimuli that underlie self-referential and other-referential action^33,72,73^.

Mothers with higher social cognition also showed reduced inhibition from the dorsomedial prefrontal cortex to the nucleus accumbens. However, this connection was only significant once accounting for wellbeing. Importantly, while the nucleus accumbens-amygdala-parahippocampal gyrus limbic network subserves *affective* theory of mind and empathy, the dorsomedial prefrontal cortex is involved in a separate but parallel *cognitive* theory of mind/empathy network^33^. This indicates that improved social cognition in motherhood is associated with reduced top-down inhibition *of* the affective theory of mind network *from* the cognitive theory of mind network. Interestingly, the link between the dorsomedial prefrontal cortex and nucleus accumbens in the group-by-cognition and group-by-social cognition models were reciprocal. In the cognition model, the information flow was top-down (dorsomedial prefrontal cortex to nucleus accumbens) and in the social cognition model, the information flow was bottom-up (nucleus accumbens to dorsomedial prefrontal cortex). Both analyses indicate that improved cognition is associated with disinhibition of the links between more cognitively-oriented regions (dorsomedial prefrontal cortex) and the reward-based and affective limbic system (nucleus accumbens-amygdala-parahippocampal gyrus). The dorsomedial prefrontal cortex-nucleus accumbens connection in both models was mediated by wellbeing (depression, anxiety and sleep), indicating that disinhibition or increased responsiveness is affected by an individual’s daily functioning. The dependence of the dorsomedial prefrontal cortex - nucleus accumbens connection on wellbeing in these models (Supplementary Information 2) is compatible with the known physiology of this connection in sleep, memory, cognition^69-71^.

### Influence of maternal caregiving on effective connectivity within the parental caregiving network

Strikingly, every node within the studied network was influenced by maternal caregiving. These regions, previously hypothesised to form a parental caregiving network^17^, do indeed appear to function together as a network related to maternal caregiving in mothers (dorsomedial prefrontal cortex was also influenced by caregiving, but only when controlling for wellbeing; see Supplementary Information 2). Specifically, the affective limbic network was particularly implicated in maternal caregiving, with reciprocal increases and decreases in inhibition with positive maternal care. Interestingly, information transfer within the affective limbic network occurred via the nucleus accumbens. Reciprocal increases and decreases in inhibition occurred between the nucleus accumbens and parahippocampal gyrus, and between the nucleus accumbens and amygdala, but not between the parahippocampal gyrus and amygdala. Models of the affective limbic system posit that the nucleus accumbens is the key ‘nexus’ or hub region within this network, and plays a major role in action selection in goal-based behaviour^73^. The nucleus accumbens integrates motivational value information from the parahippocampal gyrus and amygdala to inform goal selection, to ensure that goals with high value are selected. This system facilitates action selection so that the individual can rapidly and flexibly switch between goals, with the nucleus accumbens integrating multiple sources of information from the parahippocampal gyrus and amygdala to assign motivational value to those goals. Goals with high motivaltional salience are more likely to be selected over those with less motivational salience^73^. The outcome of the information transfer processes within the affective limbic network is ultimately the behavioural goal that the individual pursues. That is, this result suggests that positive maternal caregiving behaviour is highly dependent on information transfer within the affective limbic network, with the nucleus accumbens serving as the hub. The motivational salience of caregiving behaviour is highlighted by the fact that the amygdala and nucleus accumbens are consistently activated in response to infant cues in task-based fMRI studies^17,24,74-78^. Experiencing increased pleasure and reward in response to an infant increases the salience of infant cues, and promotes increased attention and attachment, which serves as a positive feedback loop to maintain sensitive caregiving behaviours^79,80^. Functional connectivity between the amygdala and the nucleus accumbens has previously been shown to relate to positive maternal care^3,24^. Specifically, mothers who were later into the postpartum period had increased functional connectivity between the left amygdala and the nucleus accumbens, which was associated with positive maternal behaviour, such as the ability to scaffold appropriate mother-child interactions^3^.

Inhibitory connection from the dorsomedial prefrontal cortex to the affective limbic network (this time to the amygdala) was implicated in maternal caregiving, but only when controlling for wellbeing (Supplementary Information 2). The ventromedial prefrontal cortex, which is more directly implicated in affective mentalising and theory of mind^33^ was not implicated. Intriguingly, *interaction* of the cognitive and affective mentalising networks may ne implicated in maternal caregiving, rather than specifically affective *or* cognitive aspects of the mentalising network. Top-down connection from the dorsomedial prefrontal cortex to the limbic network was more disinhibited (i.e., more excitable) with increased positive maternal caregiving, and modulated by wellbeing, suggesting that the interaction between cognitive and affective networks is most affected by wellbeing, and associated with mood disorders in the postpartum period.

While a more disinhbited network may subserve more flexible and responsive behaviour, in the extreme it may lead to hyperexcitability, and predispose the individual to pathologies like mood and anxiety disorders^81,82^. Inhibitory connections in particular are important in the pathogenesis of anxiety and fear disorders^81,83^. Activation of the amygdala and increased salience of infant cues is part of a healthy adaptation to motherhood, giving rise to increased vigilance, protectiveness, threat detection, and preoccupation with the infant^84,85^. However, the brain circuits important for adaptive ‘checking and worrying’ behaviours, overlap with those that are hyperactive in anxiety disorders^79^. Whilst plasticity in salience networks is a healthy part of threat detection and harm avoidance^86^, aberrant functional adaptations – abnormally decreased or excessive worry – may be related to postpartum psychopathology^79^. Importantly, as the participants in this study were healthy and did not meet the criteria for depression or anxiety (see Results) the results represent variability within the healthy postpartum experience.

Mothers with higher maternal caregiving also showed decreased inhibition of the posterior cingulate cortex from the ventromedial prefrontal cortex, and nucleus accumbens, indicating an important role of the posterior cingulate in positive maternal care^28^. Within the ‘dynamic systems’ model of posterior cingulate function, increased activity of the dorsal posterior cingulate allows increased whole-brain meta-stability, allowing efficient and rapid transitions between neural states, and more efficient cognition. We found that increased caregiving is associated with directional reductions of dorsal posterior cingulate inhibition from the ventromedial prefrontal cortex and nucleus accumbens, consistent with the interpretation that those with higher caregiving scores can recruit more efficient and flexible cognitive system in response to salient, behaviourally-rewarding stimuli (i.e., the infant). This result complements earlier findings in mothers with postpartum depression, whereby the posterior cingulate cortex showed reduced resting state functional connectivity with the parahippocampal gyrus, ventromedial prefrontal cortex, and amygdala^18^. Postpartum depression affects the quality of maternal caregiving^87^, further underlying the importance of ventromedial prefrontal cortex/limbic inhibitory connections with the posterior cingulate in positive maternal caregiving.

### Selective Modulation of Inhibitory Connectivity in Motherhood

Our findings suggest a selective modulation of inhibitory pathways in motherhood. Inhibition in a cortical hierarchy serves to maintain efficient neural signalling by modulation of the primary inhibitory neurotransmitter, gamma-aminobutyric acid (GABA)^60,88^. In pregnancy and the postpartum period, GABAergic activation is modulated by fluctuating concentrations of neurosteroids, such as allopregnanolone (a metabolite of progesterone). Allopregnanolone acts at the GABA-A receptor to increase inhibition^89,90^, and GABA-A receptor density and sensitivity are flexibly upregulated and downregulated to maintain an ideal homeostatic level of inhibition^88,89^ (i.e., at times of high allopregnanolone, receptor density and sensitivity are down-regulated, and vice versa). Allopregnanolone concentrations fluctuate across the peripartum period, with high levels in late pregnancy, followed by a rapid decline at parturition. Allopregnanolone levels remain low throughout the postpartum period until the return of menstruation, and cessation of breastfeeding. Weaning therefore represents an increase in the concentration of allopregnanolone and a period of GABAergic plasticity^91^. Our observed differences in inhibitory connectivity in mothers at one-year postpartum may reflect the reorganisation of the GABAergic system at this time^30^, providing a potential mechanism for our results. Plasticity of the GABAergic system across the peripartum is related to maternal behaviour, and dysregulation of GABAergic signalling in rodents results in deficits in maternal care, and increased symptoms of anxiety and depression^88^. Our interpretation of the selective modulation of inhibitory connectivity would be strengthened with the addition of hormonal data, which was not collected from this sample. Therefore, it is not possible to assert that the differences we observed are related to levels of allopregnanolone. Future research should investigate the relationships between fluctuating hormone levels and inhibitory connectivity across the peripartum period.

### Study Context & Future Directions

The study findings should be interpreted within the context of the design and limitations. The cross-sectional study design limits the results to *differences* between groups, and not *changes* across the peripartum period^92,93^. Future studies could use longitudinal samples to examine how these networks develop across the transition to motherhood. The study included a modest sample size of 40 mothers and 39 non-mothers. Whilst this is considered a large sample size for conducting spDCM analyses^94^, and in studies of parental neuroscience^2,4,6,8^, larger samples may be able to detect more subtle effects. The dimensionality of the spDCM model, while large in comparison to previous spDCM models (e.g.,^95^), is probably low with respect to the known widespread effects of parenthood on the brain (e.g.,^10,17,78^). spDCM is a relatively new but highly robust analytical approach to investigate effective connectivity between regions. As it is computationally demanding, spDCM is most suited to models with low dimensionality, and hypothesis-driven seeds. For this reason, we chose to reduce the dimensionality of the model space by examining left-lateralised regions only. We had no strong hypotheses regarding the laterality of the effects, and the studies that informed our selection of ROIs (Figure 1b) showed a slight tendency towards left-than right-lateralised results. While it is typical in network neuroimaging to reduce the model dimensionality to aid computation and interpretation of the final results, the results of spDCM are specific to the regions of interest used in the model, and studies using different seed regions may yield different effects. As more studies investigate resting state brain connectivity in motherhood, different brain regions may emerge as those most relevant for understanding motherhood and caregiving behaviour.

### Conclusions

In summary, the observed patterns of disinhibition indicate a maternal brain that is more efficient, flexible and responsive at rest. The consistency of this pattern across analyses suggests that these differences generalise across diverse behaviours in the postpartum period, including those that are related to maternal caregiving, as well as non-maternal behaviours, i.e., cognition and social cognition. Together our results support the interpretation of these key regions as constituting a parental caregiving network, with the nucleus accumbens and the parahippocampal gyrus emerging as ‘hub’ regions, highlighting the global importance of the affective limbic network for maternal caregiving, social cognition, and cognitive performance in the postpartum period.

## Methods

The study protocol was approved by Monash University Human Research Ethics Committee (Project ID: 19455) and conducted in accordance with the Australian National Statement on Ethical Conduct in Human Research (2007).

### Participants

Our sample comprised 86 women, 43 first-time mothers approximately one-year postpartum (11.7**±**1.7 months) and 43 women who had never been pregnant. All participants were aged over 18 years and right-handed. Inclusion criteria for the study included fluency in English, not currently pregnant, no personal or family history of psychiatric disorder except depression and anxiety, no history of head injury, neurological disorder, epilepsy or stroke, and completion of an MRI safety survey for subsequent MRI scanning (e.g., presence of metal in the body, past surgeries, and claustrophobia). Seven participants were excluded due to incomplete or insufficient fMRI scans: two participants had incomplete scans, one had an insufficient imaging field of view issue, and four had poor image segmentation results. Our final sample included 40 mothers and 39 non-mothers.

### Procedure

After providing informed consent, all participants completed a pregnancy test, and then completed a psychosocial and cognitive battery, including paper and computer-based tasks and questionnaires. Demographic information and medical and obstetric history (including information about pregnancy, birth, breast-feeding, and time spent in primary care) was collected via clinical interview. Cognitive measures included the Hopkins Verbal Learning Test (HVLT; verbal memory)^36^, digit span forward and backward (working memory maintenance and manipulation, respectively)^37^, Symbol-Digit Modalities Task (SDMT; processing speed)^38^; Reading the Mind in Films Task (RMFT; Theory of Mind)^39^, Toronto Empathy Questionnaire^40^, and the Prospective and Retrospective Memory Questionnaire (PRMQ; subjective memory)^41^. Psychosocial measures included the Pittsburgh Sleep Quality Index (PSQI)^42^, Beck Depression Inventory (BDI)^43^, Beck Anxiety Inventory (BAI)^44^, Maternal Postnatal Attachment Scale (MPAS)^45^, and Maternal Self-Efficacy Questionnaire (MEQ)^46^. Participants completed the PRMQ, BDI, BAI, MPAS, and MEQ questionnaires at a time of their convenience on their own devices (e.g., laptop, phone) via a Qualtrics survey, before attending their testing session. The relationships between cognitive performance, self-perception of memory ability, and wellbeing (sleep, depression and anxiety) in this sample are discussed in detail in Orchard et al.^35^.

### MRI Acquisition

All MRI scans were obtained on a Siemens 3T Skyra MR scanner (Erlangen, Germany) with a 32-channel head and neck coil at Monash Biomedical Imaging, Melbourne, Australia. T1-weighted magnetization-prepared rapid gradient-echo (MP2RAGE) images were acquired (TR=5000ms, TE=2.98ms, TI 1=700ms, TI 2=2500ms, matrix size=256×240×172, bandwidth=240 Hz/pixel) and used for anatomical reference. T2-weighted images (TR=3200ms, TE=452ms, matrix size=206×206×128, bandwidth=698 Hz/pixel) were also used for anatomical reference. Resting state fMRI (rsfMRI) data were obtained using a multiband, multi-echo planar imaging (EPI) sequence (TR=910ms; multiband acceleration factor=4; matrix size=64×64; number of slices=40; TE 1=12.60ms; TE 2=29.23ms; TE 3=45.86; TE 4=62.49; number of time points=767; bandwidth=2520 Hz/pixel; 3.2mm isotropic voxel). Following functional data acquisition, a reverse-phase scan was acquired to correct the rsfMRI scan for geometric distortions.

### MRI processing and denoising

All images were first processed using a standardised pipeline, *fmriprep*^47^. Briefly, the pipeline included slice time correction, susceptibility distortion correction using gradient field-maps, co-registration to the corresponding T1w using boundary-based registration with six degrees of freedom. Motion correcting transformations, BOLD-to-T1w transformation and T1w-to-template (MNI) warp were concatenated and applied in a single step. No participants exceeded a previously established head motion exclusion criteria^48^ (mean framewise displacement>.55mm). For each scan, we then applied an automated TE-dependent ICA-based multi-echo de-noising pipeline, *tedana*^49^, resulting in an optimally combined fMRI image^50^. Images were further denoised by regressing out averaged signals from the white matter and cerebral spinal fluid voxel wise, prior to high-pass filtering (f>.005 Hz) and linear detrending. Quality control metrics, functional connectivity matrices and carpet plots were visualised to ensure the pre-processing and denoising steps achieved the desired effects of reducing noise and association between head motion and functional connectivity^48^. The full quality control protocol and report can be found at: https://github.com/sidchop/NAPPY_DCM.

### Data Analysis

#### Behavioural Data

Full behavioural methods and results are reported in Orchard, et al. ^35^. Briefly, we used principal components analyses (PCA) to compute summary scores using z-scored measures of cognitive performance (Hopkins Verbal Learning Test, Digit Span Forward, Digit Span Backward, and Symbol Digit Modalities Task), social cognition (Reading the Mind in the Films Task, Toronto Empathy Questionnaire), maternal caregiving (Maternal Attachment Questionnaire and Maternal Self-Efficacy Questionnaire), and wellbeing (Pittsburgh Sleep Quality Index, Beck Depression Inventory and Beck Anxiety Inventory). PCs with eigenvalues >1 were retained^51^ for further use within the spectral dynamic causal modelling (spDCM) analyses.

#### Regions of Interest

Six regions of interest (ROIs) were identified as key nodes for subsequent spDCM analysis based on the available functional and effective connectivity literature in motherhood (Figure 1) ^18,22,24,30^, and compared against the relevant Brodmann areas mapped in MNI space to ensure consistency between region labelling^52^. The identified neural network comprised the left dorsomedial prefrontal cortex, ventromedial prefrontal cortex, posterior cingulate cortex, parahippocampal gyrus, amygdala, and nucleus accumbens (Figure 1a). As spDCM is computationally demanding, it is most suited to models with low dimensionality and hypothesis-driven seeds. Furthermore, there was no evidence to suggest laterality of the effects and studies that informed our selection of ROIs (Figure 1b) showed a slight tendency towards left-than right-lateralised results. We therefore examined only left-lateralised regions to reduce the model space. To extract BOLD time series corresponding to the six ROIs, we selected the MNI coordinates as the centre of a 5-mm (dorsomedial prefrontal cortex, ventromedial prefrontal cortex, posterior cingulate cortex, and parahippocampal gyrus) or 3-mm sphere (amygdala and nucleus accumbens), which were then used to compute the subject-specific principal eigenvariate and correct for confounds.

#### Neural network modelling: Spectral Dynamic Causal Modelling (spDCM)

The spDCM analyses were performed using the functions of DCM12.5 (revision 7497) implemented in SPM12 (Supplementary Information 1). In order to assess our main hypothesis, we focused on spDCM analyses that assessed differences in the effective connectivity within the parental caregiving network (1) for mothers versus non-mothers, (2) modulated by cognition and (3) social cognition in mothers versus non-mothers, (4) modulated by maternal caregiving in mothers.

At the first-level, a fully-connected model was created for each participant. Using spDCM, we estimated the DCMs, fitted the complex cross-spectral density using a parameterised power-law model of endogenous neural fluctuations^31^. The analysis allowed for the measurement of causal interactions between regions, as well as the amplitude and exponent of endogenous neural fluctuations within each region^31^. Model inversion was based on standard variational Laplace procedures^53^. This method of Bayesian inference uses Free Energy as a proxy for (log) model evidence, while optimising the posterior density under a Laplace approximation of model parameters.

To characterise how group differences in neural circuitry were modulated by maternal caregiving, cognition, and social cognition, hierarchical models over the parameters were specified within a hierarchical Parametric Empirical Bayes (PEB) framework for DCM^54^. We based these models on our hypotheses. For each model, all continuous behavioural regressors (cognition, social cognition, and maternal caregiving) were mean centred so that the intercept of each model was interpretable as the group mean connectivity^55,56^. Group factor was modelled as the main regressor of interest as a vector consisting of 1 (mothers) and -1 (non-mothers). The interaction terms were created by first centring the continuous variables (cognition, social cognition) and then creating an element-by-element product of the newly centred variables with the categorical grouping variable. Maternal caregiving scores were modelled as main regressor of interest, whereas the remaining scores were modelled as regressors of no interest in all models. Bayesian model reduction was used to test all reduced models within each parent PEB model, assuming that a different combination of connections could exist^54^ and ‘pruning’ redundant model parameters. Parameters of the best pruned models in the last Occam’s window were averaged and weighted by their evidence (i.e., Bayesian Model Averaging) to generate final estimates of connection parameters. To identify important effects, i.e., changes in directed connectivity, we compared models using log Bayesian model evidence to ensure the optimal balance between model complexity and accuracy. Models were compared with and without each effect and the posterior probability was calculated for each model using the softmax function of the log Bayes factor. We treat effects (i.e., connection strengths and their changes) with a posterior probability >0.95 (i.e., strong evidence) as significant for reporting purposes.

##### Hypothesis 1

To investigate the effect of mothers versus non-mothers on the intrinsic effective connectivity of the parental caregiving network, the following hypothesis was tested within the PEB framework:

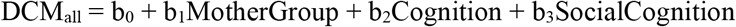

##### Hypothesis 2

To investigate the interaction effect between the mother group and cognition, the following hypothesis was tested within the PEB framework:

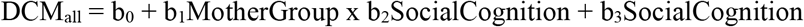

##### Hypothesis 3

To investigate the interaction effect between the mother group and social cognition, the following hypothesis was tested within the PEB framework:

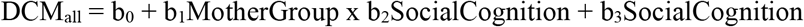

##### Hypothesis 4

To investigate the effect of maternal caregiving in mothers on the intrinsic effective connectivity of the parental caregiving network, the following hypothesis was tested within the PEB framework:

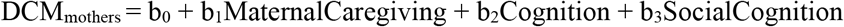

Finally, following our earlier result^35^ where we found that differences between mothers and non-mothers in cognition were accounted for by differences in wellbeing, we hypothesised that connectivity in the four comparisons will be affected by poorer *wellbeing* (sleep, depression, and anxiety). In general, the effect of wellbeing on effective connectivity was modest, and results are provided in Supplementary Information 2.

## Acknowledgements

The authors are supported by the Australian Research Council (ARC) Centre of Excellence for Integrative Brain Function (CE140100007), and this study received support from an early career researcher project grant from the Centre to ERO. SDJ is supported by an Australian National Health and Medical Research Council Fellowship (APP1174164).

## Author contributions

The study was conceptualised by ERO, SDJ, PGDW & GFE. Study funding was sourced by ERO & SDJ. Data was collected by ERO. Data was analysed by ERO, KV, SC & TT. The first draft of the manuscript was written by ERO, SDJ & KV with edits by SC, TT, PGDW & GFE. All authors have read and approved the final version of the manuscript.

### Data Availability Statement

All data supporting the results of this study are presented in the tables and supplements of the article. For access to raw data please contact the corresponding author.

