## Supplementary Information for "The maternal brain is more flexible and responsive at rest: effective connectivity of the parental caregiving network in postpartum mothers"

**1. Spectral Dynamic Causal Modelling**

Dynamic causal modelling (DCM) is a Bayesian framework that allows directed (causal) connections to be inferred between neural systems: known as *effective connectivity*. In resting-state data, the DCM approach is based on a deterministic model that generates cross-spectra predictions, and is referred to as *spectral DCM* (spDCM)^1^. We have previously applied spDCM to studies of resting-state connectivity in obesity^2^ and decision-making^3^. Full details of the spDCM approach is provided in Friston et al.^1^ and Voigt et al. ^2^, but is described briefly here.

In order to model brain activity in the absence of external stimuli or evoked responses, the spDCM model adds a stochastic component to the classic DCM based on ordinary differential equations. Mathematically, the stochastic generative model is expressed using two equations:

The neuronal state equation (S1):

$\dot{x}\left( t \right)=f\left( x(t),u(t),\theta\right)+ v(t)$	(S1)

And the observation equation (S2), a static nonlinear mapping from the hidden physiological states in (S1) to the observed BOLD activity:

$y\left( t \right)=h\left( x(t),\varphi\right)+ e\left( t \right),$	(S2)

where x ̇(t) is the rate of change of the neuronal states x(t), θ are unknown parameters (i.e., the effective connectivity) and v(t) (resp. e(t)) is the stochastic process – called the state noise (resp. the measurement or observation noise) – modelling the random neuronal fluctuations that drive the resting state activity. In the observation equations, φ are the unknown parameters of the (haemodynamic) observation function and u(t) represents any exogenous (or experimental) inputs that drive the hidden states – that are usually absent in resting state designs. spDCM furnishes a constrained inversion of the stochastic model by parameterising the neuronal fluctuations$v(t)$. spDCM simplifies the generative model by replacing the original timeseries with its second-order statistic, the cross-spectra. As such, instead of estimating time-variant hidden states, spDCM estimates their temporally-invariant covariance.

The parametric empirical Bayes (PEB) framework^4^ was used to test for differences in effective connectivity. Empirical Bayes refers to the process of Bayesian model inversion or fitting of hierarchical models. In a hierarchical model, the constraints on the posterior density of model parameters at a given level is provided by the level above. The constraints are called empirical priors because they are informed by the data. PEB is a second-level model over parameters, and represents how individual within-subject connections derive from the subjects’ group membership. PEB relies upon Bayesian Model Reduction (BMR) to refine the inversion of multiple models of a single dataset, or a single hierarchical model of multiple datasets. BMR can improve subject-specific parameter estimates by using group-level estimates to inform the calculation of individual DCMs from local optima.

**2. Effect of Wellbeing**

In general, the effect of wellbeing was modest across all analyses (Supplementary Figure 1). For the main effect of group, no additional connections were identified when controlling for wellbeing (Supplementary Table 6). In the cognition analysis, when controlling for the effect of wellbeing, mothers with better cognitive performance show increased self-inhibition of the parahippocampal gyrus, but no longer show the bottom-up inhibition from the nucleus accumbens to the dorsomedial prefrontal cortex (Supplementary Table 7). For social cognition, mothers with better social cognition also showed decreased top-down inhibition of the nucleus accumbens from the dorsomedial prefrontal cortex when controlling for the effect of wellbeing (Supplementary Table 8). Finally, for maternal caregiving, mothers with better maternal caregiving also showed increased inhibition of the amygdala from the nucleus accumbens, and decreased top-down inhibition of the amygdala from the dorsomedial prefrontal cortex when controlling for the effect of wellbeing (Supplementary Table 9).

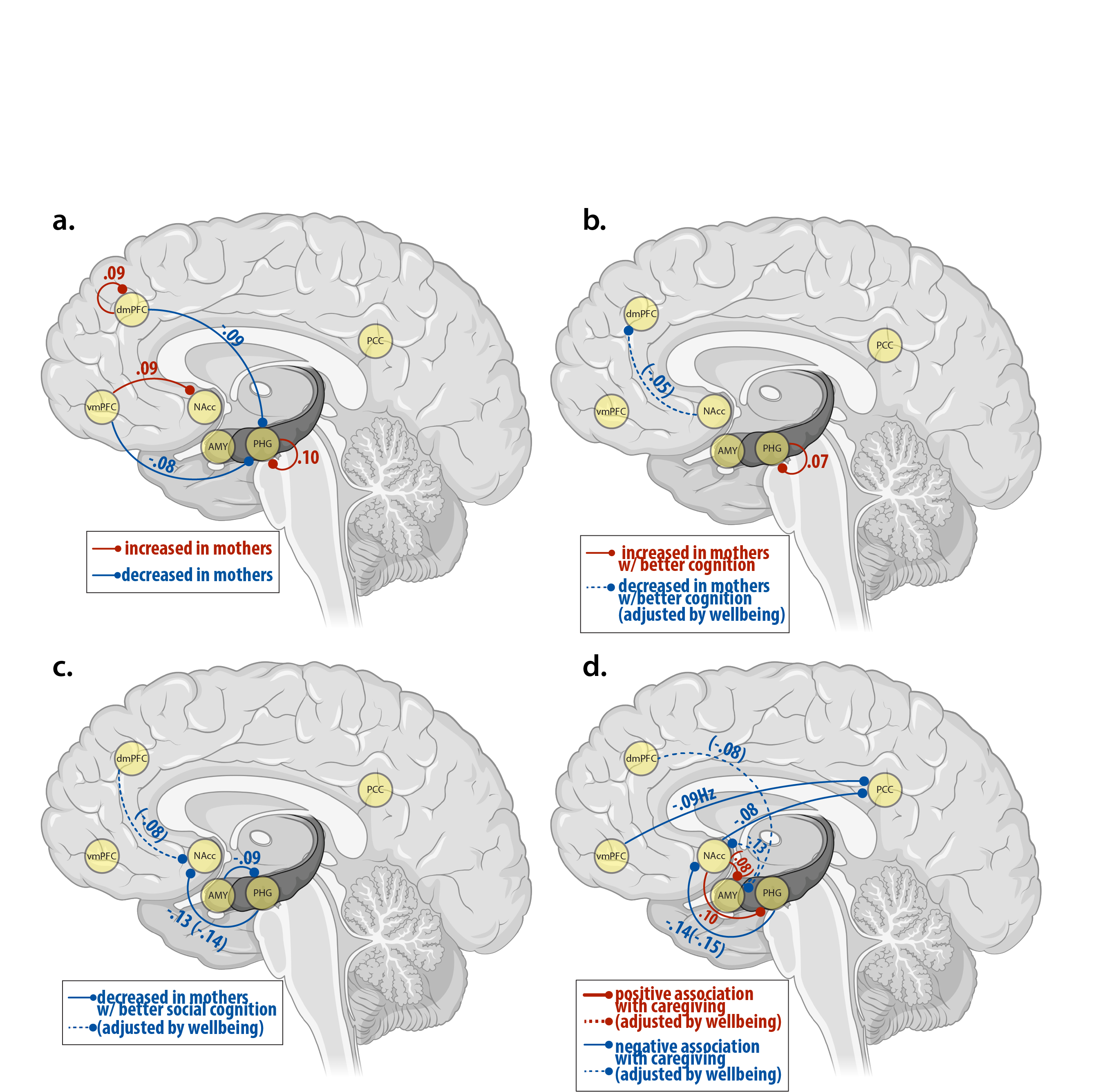

**Supplementary Figure 1:** *Schematic diagram showing spDCM results for (a.) the main effect of group (mothers vs non-mothers), (b.) the interaction effect of group-by-cognition, (c.) the interaction effect of group-by-social cognition, and (d.) the main effect of maternal caregiving in mothers only. All connections are inhibitory. Values represent effect sizes (see Table 2-5) in Hz. Blue lines depict decreased inhibition and red lines depict increased inhibition. Dashed lines represent connections that are attenuated when the model adjusts for wellbeing, and solid lines are connections which are present whether or not wellbeing is adjusted for. Abbreviations: PCC, posterior cingulate cortex; vmPFC, ventromedial prefrontal cortex; dmPFC, dorsomedial prefrontal cortex; PHG, parahippocampal gyrus; Amyg, amygdala; NAcc, nucleus accumbens.*

| **Supplementary Table 6:** *Effect size, valence and posterior probabilities for each connection showing a main effect of group (mothers>non-mothers)* | | | | | |
| --- | --- | --- | --- | --- | --- |
| **Connection** | **Valence** | ***Mothers vs. Non-Mothers*** | | ***Mothers vs. Non-Mothers***  ***(controlled by wellbeing)*** | |
|  |  | Effect Size in Hz [95% CI] | Posterior Probability | Effect Size in Hz  [95% CI] | Posterior Probability |
| dmPFC🡪 dmPFC | Inhibition | 0.09 [0,0.16] | 0.97 | 0.08 [0, 0.16] | 0.96 |
| dmPFC 🡪 PHG | Inhibition | -0.09 [-0.19,0] | 0.97 | -0.10 [-0.19, -0.02] | 0.98 |
| vmPFC 🡪 PHG | Inhibition | -0.08 [-0.16,0] | 096 | -0.08 [-0.16, 0] | 0.95 |
| vmPFC 🡪 NAcc | Inhibition | 0.09 [0.01,0.17 | 0.96 | 0.09 [0.01, 0.17] | 0.96 |
| PHG 🡪 PHG | Inhibition | 0.10 [0.18,0.01] | 0.98 | 0.11 [0.03, 0.19] | 0.99 |
| *Abbreviations: Hertz (Hz), Confidence Interval (CI), dorsomedial prefrontal cortex (dmPFC), ventromedial prefrontal cortex (vmPFC), parahippocampal gyrus (PHG), amygdala (Amyg) nucleus accumbens (NAcc).* | | | | | |

| **Supplementary Table 7:** *Effect size, valence and posterior probabilities for each connection showing an interaction effect of group-by-cognition, with and without controlling for wellbeing* | | | | | |
| --- | --- | --- | --- | --- | --- |
| **Connection** | **Valence** | ***Group-by-Cognition*** | | ***Group-by-Cognition***  ***(controlling for Wellbeing)*** | |
|  |  | Effect Size in Hz [95% CI] | Posterior Probability | Effect Size in Hz  [95% CI] | Posterior Probability |
| PHG 🡪 PHG | Inhibition | 0.07 [0.01, 0.12] | 0.98 | 0.07 [0.015, 0.126] | 0.98 |
| NAcc 🡪 dmPFC | Inhibition | -0.05 [-0.11, -0.00] | 0.95 |  |  |
| *Abbreviations: Hertz (Hz), Confidence Interval (CI), dorsomedial prefrontal cortex (dmPFC), parahippocampal gyrus (PHG), nucleus accumbens (NAcc).* | | | | | |

| **Supplementary Table 8:** *Effect size, valence and posterior probabilities for each connection showing an interaction effect of group-by-social cognition, with and without controlling for wellbeing* | | | | | |
| --- | --- | --- | --- | --- | --- |
| **Connection** | **Valence** | ***Group-by-Social Cognition*** | | ***Group-by-Social Cognition***  ***(controlling for Wellbeing)*** | |
|  |  | Effect Size in Hz [95% CI] | Posterior Probability | Effect Size in Hz  [95% CI] | Posterior Probability |
| PHG 🡪 NAcc | Inhibition | -0.13 [-0.21, -0.05] | 1.00 | -0.14 [-0.22, -0.05] | 1.00 |
| Amyg 🡪 PHG | Inhibition | -0.09 [-0.18, -0.01] | 0.97 | -0.09 [-0.18, -0.01] | 0.97 |
| dmPFC 🡪 NAcc | Inhibition |  |  | -0.08 [-0.16, -0.00] | 0.95 |
| *Abbreviations: Hertz (Hz), Confidence Interval (CI), dorsomedial prefrontal cortex (dmPFC), ventromedial prefrontal cortex (vmPFC), parahippocampal gyrus (PHG), amygdala (Amyg) nucleus accumbens (NAcc).* | | | | | |

| **Supplementary Table 9:** *Main effect of maternal caregiving, with and without controlling for wellbeing* | | | | | |
| --- | --- | --- | --- | --- | --- |
| **Connection** | **Valence** | ***Maternal Caregiving*** | | ***Maternal Caregiving***  ***(controlling for Wellbeing)*** | |
|  |  | Effect Size in Hz [95% CI] | Posterior Probability | Effect Size in Hz  [95% CI] | Posterior Probability |
| vmPFC 🡪 PCC | Inhibition | -0.09 [-0.17, -0.01] | 0.97 | -0.09 [-0.02, -0.02] | 0.98 |
| PHG 🡪 NAcc | Inhibition | -0.14 [-0.22, -0.06] | 1.00 | -0.15 [-0.22, -0.07] | 1.00 |
| Amyg 🡪 NAcc | Inhibition | -0.13 [-0.20, -0.05] | 1.00 | -0.13 [-0.20, -0.05] | 1.00 |
| NAcc 🡪 PCC | Inhibition | -0.08 [-0.16, -0.01] | 0.97 | -0.08 [-0.16, -0.00] | 0.96 |
| NAcc 🡪 PHG | Inhibition | 0.1 [0.02, 0.18] | 0.98 | 0.1 [0.03, 0.18] | 0.99 |
| dmPFC 🡪 Amyg | Inhibition |  |  | -0.08 [-0.16, -0.00] | 0.96 |
| NAcc 🡪 Amyg | Inhibition |  |  | 0.08 [0.00, 0.15] | 0.95 |
| *Abbreviations: Hertz (Hz), Confidence Interval (CI), dorsomedial prefrontal cortex (dmPFC), ventromedial prefrontal cortex (vmPFC), parahippocampal gyrus (PHG), posterior cingulate cortex (PCC), amygdala (Amyg), nucleus accumbens (NAcc).* | | | | | |

**3. Cognition Summary Scores**

| **Supplementary Table 1:** *Cognitive Performance* *Principal Component Coefficient Loadings, Variance Explained, and Eigenvalues for each component.* | | | | | |
| --- | --- | --- | --- | --- | --- |
|  | **PC1** | | **PC2** | **PC3** | **PC4** |
| Hopkins Verbal Learning Test | 0.48 | | 0.48 | 0.73 | -0.06 |
| Digit Span Forwards | 0.54 | | -0.43 | -0.02 | 0.72 |
| Digit Span Backwards | 0.53 | | -0.49 | -0.07 | -0.69 |
| Symbol Digit Modalities Task | 0.44 | | 0.59 | -0.68 | -0.00 |
| Variance Explained (%) | 54.59 | | 23.18 | 13.32 | 8.91 |
| Eigenvalue | **2.18** | | 0.93 | 0.53 | 0.36 |
| **Supplementary Table 2:** *Social Cognition* *Principal Component Coefficient Loadings, Variance Explained, and Eigenvalues for each component.* | | | | | |
|  | | **PC1** | | | **PC2** |
| Reading the Mind in the Films Task | | 0.71 | | | 0.71 |
| Toronto Empathy Questionnaire | | 0.71 | | | -0.71 |
| Variance Explained (%) | | 51.20 | | | 46.76 |
| Eigenvalue | | **1.02** | | | 0.94 |

| **Supplementary Table 3:** *Maternal Caregiving* *Principal Component Coefficient Loadings, Variance Explained, and Eigenvalues for each component.* | | |
| --- | --- | --- |
|  | **PC1** | **PC2** |
| Maternal Postnatal Attachment Questionnaire | 0.71 | -0.71 |
| Maternal Self-Efficacy Questionnaire | 0.71 | 0.71 |
| Variance Explained (%) | 84.22 | 16.81 |
| Eigenvalue | **1.68** | 0.34 |

| **Supplementary Table 4:** *Wellbeing* *Principal Component Coefficient Loadings, Variance Explained, and Eigenvalues for each component.* | | | |
| --- | --- | --- | --- |
|  | **PC1** | **PC2** | **PC3** |
| Pittsburgh Sleep Quality Index | 0.55 | 0.82 | -0.16 |
| Beck Depression Inventory | 0.60 | -0.26 | 0.76 |
| Beck Anxiety Inventory | 0.59 | -0.51 | -0.63 |
| Variance Explained (%) | 66.73 | 19.25 | 14.02 |
| Eigenvalue | **2.00** | 0.58 | 0.42 |

| **Supplementary Table 5:** *Means and standard deviations of tasks and questionnaires which comprise the cognitive performance, social cognition and wellbeing principal components.* | | |
| --- | --- | --- |
|  | **Mean (SD)**  Mothers | **Mean (SD)**  Non-Mothers |
| Cognitive Performance |  |  |
| *Hopkins Verbal Learning Test* | 31.18 (3.1) | 30.64 (3.8) |
| *Digit Span Forward* | 7.03 (1.2) | 7.59 (1.2) |
| *Digit Span Backward* | 6.43 (1.0) | 6.87 (1.4) |
| *Symbol Digit Modalities Task* | 58.73 (7.3) | 61.64 (9.9) |
| Social Cognition |  |  |
| *Reading the Mind in the Films Task* | 66.59 (9.2) | 63.17 (11.1) |
| *Toronto Empathy Questionnaire* | 50.18 (4.8) | 46.70 (4.2) |
| Wellbeing |  |  |
| *Pittsburgh Sleep Quality Index* | 7.33 (3.1) | 4.67 (2.6) |
| *Beck Depression Inventory* | 8.55 (5.6) | 4.95 (4.3) |
| *Beck Anxiety Inventory* | 10.70 (8.6) | 8.56 (6.63) |

**4. Nucleus Accumbens and Parahippocampal Gyrus: Hub Regions of the Maternal Brain**

The results indicated that the nucleus accumbens and the parahippocampal gyrus are ‘hub’ regions of the maternal caregiving brain network. Both regions showed significant connections in every model tested: main effect of group, main effect of maternal caregiving, interaction effect of group-by-cognition, and interaction effect of group-by-social cognition. Both the nucleus accumbens and parahippocampal gyrus are part of both the ventromedial emotion regulation system^5^ and the affective limbic network^6^, suggesting that the increased influence of these regions supports, and perhaps integrates, the functions of both networks.

The nucleus accumbens, part of the striatum and mesolimbic reward circuit, is considered a crucial region in the human maternal brain^7^, showing both structural and functional neural adaptations across the peripartum period. The nucleus accumbens decreases in volume up to 25% across pregnancy^8^, and at two-months postpartum^9^, with greater volume reductions associated with increased activation in response to infant stimuli^10^. The striatum also shows grey matter volume increases later in the postpartum period. Specifically, consistent increases in grey matter volume were found in the caudate^11,12^, putamen^11^, nucleus accumbens^13^, and pallidum^11^. Studies of resting state functional connectivity have also highlighted the role of the nucleus accumbens in sensitive maternal caregiving behaviours^14,15^. Increased connectivity between the nucleus accumbens and the amygdala is associated with a mother’s ability to scaffold appropriate interactions with her child^14^. Positive maternal caregiving behaviours were also related to functional connectivity between the nucleus accumbens and the dorsomedial prefrontal cortex^15^, and activation of the nucleus accumbens is correlated with increased oxytocin levels^15^. Our results, alongside the extant literature highlight the importance of the nucleus accumbens in the global functioning of the maternal brain. Our results emphasise the influence of the nucleus accumbens for maternal cognition, and social cognition, and that the increased inhibitory influence of the nucleus accumbens may play a central role in maternal caregiving.

The parahippocampal gyrus also undergoes structural plasticity in motherhood, with grey matter changes across pregnancy^8^, the postpartum period^11^, and in late-life^16^. The parahippocampal gyrus is involved in memory consolidation and cohesion^17^ and mediates cortico-hippocampal communication^18^. Altered parahippocampal structure and function is potentially related to changes in maternal cognition across pregnancy^19^. However, the involvement of the parahippocampal gyrus appears to extend beyond the cognitive domain. The parahippocampal gyrus is implicated across many studies of maternal brain function and sensitive caregiving^8,11,16^, suggesting this region has a broader role in maternal caregiving. A recent meta-analysis has highlighted the parahippocampal gyrus as one of only six brain regions that show consistent activation in response to visual stimuli of a mother’s own child, compared to an unknown control child^20^. Studies of resting state functional connectivity show disrupted connectivity between the parahippocampal gyrus, and the posterior cingulate cortex^21^, amygdala^21^, and dorsomedial prefrontal cortex^22^. Taken together, the parahippocampal gyrus is a key region of the maternal brain, which deserves increased attention in future studies.

**5. Social Cognition**

Compared to non-mothers, mothers had significantly higher scores on measures of social cognition (theory of mind, and empathy; Supplementary Tables 1; 5). Social cognition abilities contribute to successful social interactions and are important for human survival and successful maternal behaviours^23^. Motherhood requires a heightened focus on infant-related responsibilities, and for a mother to be more in tune with the thoughts, feelings, and needs of their children, especially pre-verbal infants, where infant needs must be interpreted without verbal communication^24^. This result is consistent with superior social cognition found in pregnant women, where mothers show improved ability to encode emotional faces during late pregnancy, compared to early pregnancy^25^, and increased facial recognition compared to non-pregnant women^26^. Interestingly, both the Reading the Mind in the Films Task and the Toronto Empathy Questionnaire are designed to assess social cognition in adults, are not specific to motherhood or the peripartum period, and do not contain questions or items about caring for children^27,28^. These observations support the interpretation that mothers’ superior social cognition performance may generalise beyond the context of childcare-related skills and behaviours, to abilities that may benefit mothers in their social interactions with other adults.
